# Mobile introns shape the genetic diversity of their host genes

**DOI:** 10.1101/092585

**Authors:** Jelena Repar, Tobias Warnecke

**Author notes:** Corresponding author Contact information: Tobias Warnecke, MRC Clinical Sciences Centre, Hammersmith Hospital Campus, Du Cane Road, London W12 0NN, United Kingdom.

## Abstract

Self-splicing introns populate several highly conserved protein-coding genes in fungal and plant mitochondria. In fungi, many of these introns have retained their ability to spread to intron-free target sites, often assisted by intron-encoded endonucleases that initiate the homing process. Here, leveraging population genomic data from *Saccharomyces cerevisiae, Schizosaccharomyces pombe*, and *Lachancea kluyveri*, we expose non-random patterns of genetic diversity in exons that border self-splicing introns. In particular, we show that, in all three species, the density of single nucleotide polymorphisms increases as one approaches a mobile intron. Through multiple lines of evidence we rule out relaxed purifying selection as the cause of uneven nucleotide diversity. Instead, our findings implicate intron mobility as a direct driver of host gene diversity. We discuss two mechanistic scenarios that are consistent with the data: either endonuclease activity and subsequent error-prone repair have left a mutational footprint on the insertion environment of mobile introns or non-random patterns of genetic diversity are caused by exonic co-conversion, which occurs when introns spread to empty target sites via homologous recombination. Importantly, however, we show that exonic co-conversion can only explain diversity gradients near intron-exon boundaries if the conversion templates comes from outside the population. In other words, there must be pervasive and ongoing horizontal gene transfer of self-splicing introns into extant fungal populations.

## Introduction

Self-splicing introns are selfish elements with a broad but patchy phylogenetic distribution. Found in tRNA, rRNA and (occasionally) protein-coding genes in bacteria and archaea, they are particularly numerous in mitochondrial genomes of fungi and plants where they have invaded genes encoding components of the electron transport chain (ETC) (Lambowitz and Belfort 1993). In many instances, fungal self-splicing introns have remained mobile, as demonstrated by experiments that track invasion capacity by crossing intron-containing and intron-free yeast strains (Jacquier and Dujon 1985; Wenzlau *et al.* 1989; Paschke *et al*. 1994; Lazowska *et al*. 1994) and intron presence/absence polymorphisms across natural populations of *S. cerevisiae* (Wolters *et al*. 2015), *S. pombe* (Zimmer *et al*. 1987) and *L. kluyveri* (Jung *et al*. 2012). For the majority of self-splicing introns in *S. cerevisiae*, spreading to an intron-free location is initiated by a homing endonuclease that is encoded in the intron itself and binds a large (~20-30bp), often singular target motif with high affinity [e.g. (Jacquier and Dujon 1985; Moran *et al*. 1992)]. Following cleavage of the intron-free homing site, the intron-containing copy of the mitochondrial genome is used as a template for homologous recombination (HR), resulting in the conversion of an intron-free to an intron-containing locus. Once gained, introns can be lost again either through fortuitous deletion or perhaps through a gene conversion event that involves an intronless cDNA produced by reverse transcriptase (RT) activity (Levra-Juillet *et al*. 1989), as proposed for spliceosomal introns (Cohen *et al*. 2012). In the absence of selection for intron retention, cycles of intron gain and loss ensue (Goddard and Burt 1999), accompanied by recurrent endonuclease activity that predictably targets the very same recognition site.

Here, prompted by reports of possible mutational hotspots in the vicinity of self-splicing intron (Hensgens *et al*. 1983; Zimmer *et al*. 1987; Foury *et al*. 1998), we consider what impact these invasion-loss cycles have on the genetic diversity of the host gene. In particular, we consider the possibility that endonuclease-mediated cleavage and subsequent repair might be mutagenic. Although HR is generally considered to be high-fidelity, it can carry non-negligible mutagenic risks depending on the precise nature of the repair process and whether error-prone polymerases are involved in DNA re-synthesis (Rodgers and McVey 2016). Pertinently, Hicks and colleagues observed increased mutation rates during double strand break (DSB) repair at the mating type (MAT) locus of *S. cerevisiae*, which is cleaved by the endonuclease HO and subsequently repaired via HR (Hicks *et al*. 2010).

Mutagenic side effects associated with endonuclease activity have also come into sharp focus recently with the widespread adoption of targetable endonucleases for genome engineering. The principal concern here has been to identify and reduce *off-target* activity (Cho et al. 2014; Kleinstiver et al. 2016). However, endonuclease activity can also have undesired *on-target* effects. Notably, non-homologous end joining (NHEJ) downstream of Cas9-mediated cleavage is associated with an increased risk of indel formation (van Overbeek et al. 2016). This has prompted the development of Cas9 derivatives that nick rather than cleave DNA (Cong et al. 2013; Mali et al. 2013), shifting repair pathway choice away from NHEJ and towards HR.

We reasoned that one way to develop a greater understanding of such on-target mutagenicity would be to study endonucleases in their native genomic context. If endonuclease activity is indeed mutagenic, cleavage and repair might have left a detectable imprint on population-wide genetic variation around the cleavage site. In search of such an imprint, we survey recent high-quality population genomic data from *S. cerevisiae, S. pombe*, and *L. kluyveri* to characterize single nucleotide polymorphism (SNP) patterns in exons flanking mitochondrial self-splicing introns.

## Results and Discussion

### Elevated polymorphism density at the exonic boundaries of mitochondrial introns

The *S. cerevisiae* mitochondrial reference genome harbours a single group I intron in the 21S rRNA gene and multiple group I and II introns in the protein-coding genes *cob* and *cox1* (Fig 1A). Whereas strong constraints on RNA structure and base-pairing govern the evolution of tRNA and rRNA genes throughout most of their sequence, protein-coding genes contain synonymous sites that might in principle allow for a better mutational read-out, particularly at short evolutionary time scales. We therefore focussed our analysis on protein-coding genes. Using high-coverage genome assemblies of 92 *S. cerevisiae* strains, we first considered SNP density as a function of distance from the nearest intron-exon boundary across *cox1* and *cob* exons (see Methods). We observe a marked increase in exonic SNP density as one approaches the intron-exon boundary (Kendall’s τ=−0.62, P=4.6E-14, Fig 1B), consistent with previous reports of polymorphism clusters located at the exonic border of specific endonuclease-encoding introns (Hensgens *et al*. 1983; Foury *et al*. 1998). The 5’ end of *cox1* exon 6, previously proposed as a mutational hotspot (Foury *et al*. 1998), contributes to but does not chiefly drive this trend (τ=−0.59, P=1.2E-12 when *cox1* exon 6 is excluded).

**Figure 1.**
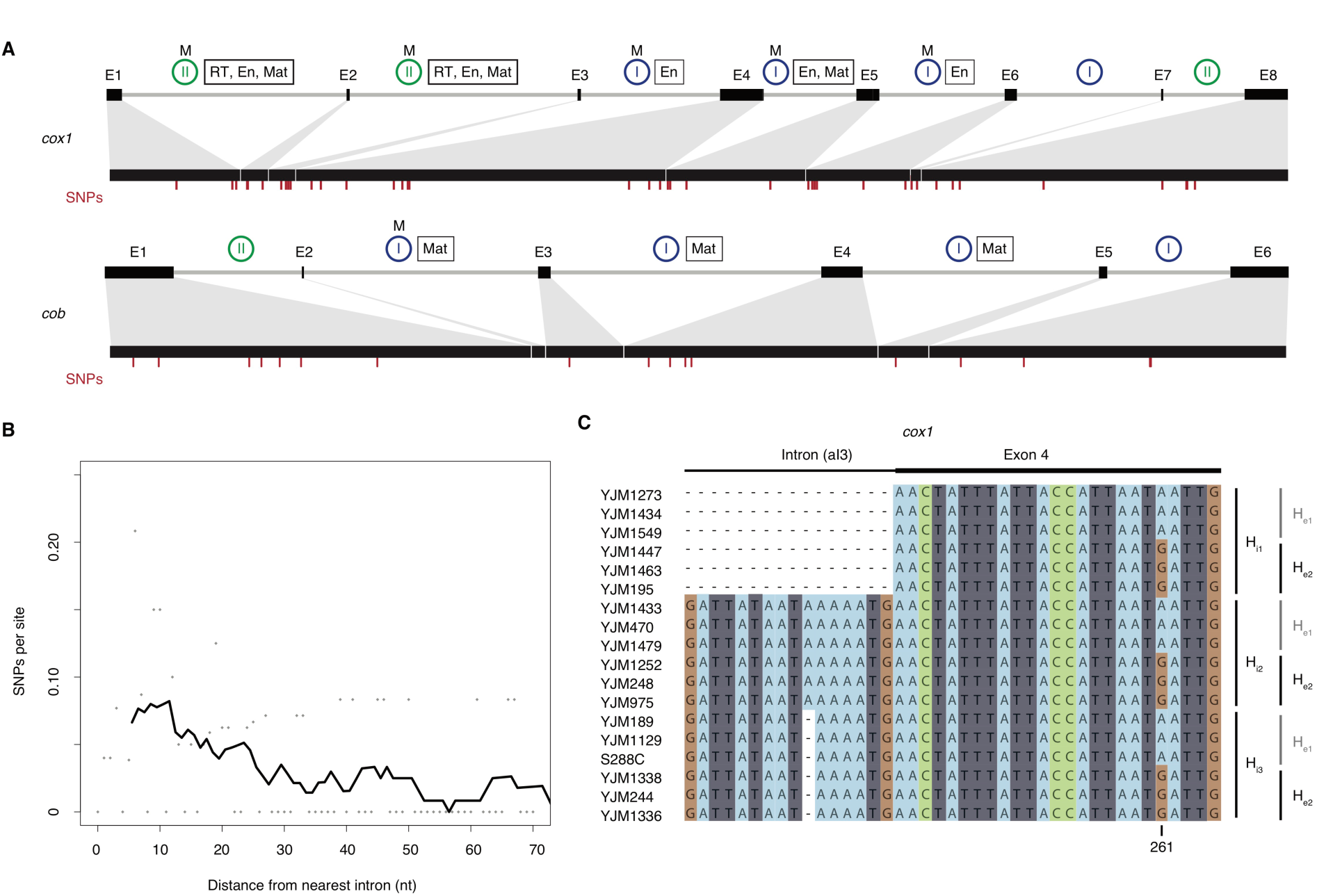
Elevated SNP density at mitochondrial intron-exon boundaries in *S. cerevisiae*. A Exon-intron structure of *cox1* and *cob*, with exons depicted as black boxes connected by grey lines (introns). Introns known to be mobile (see Methods) are labelled (M) above blue or green circles that indicate group I and II introns, respectively. Introns labelled with a rectangle harbour open reading frames that encode proteins with endonuclease (En), maturase (Mat), and/or reverse transcriptase (RT) activity. The locations of single nucleotide polymorphisms are marked by red dashes. B SNP density across all *cox1/cob* exons as function of their distance from the nearest intron-exon boundary. The density curve is calculated by smoothing across 10nt windows, moving in 1nt steps. The curve starts at centre point of first window rather than at 0. C Excerpt from the *cox1* alignment of 92 *S. cerevisiae* strains, highlighting a short region at the junction of intron 3 and exon 4 across 18 strains. Three different intronic (H_i1-3_) and two different exonic (H_e1-2_) haplotypes are evident, with all six possible combinations present in the population.

In addition, the SNP density gradient is evident for both group I introns (τ=−0.51, P=6.4E-10) and group II introns (τ=−0.60, P=6.4E-12) and is broadly similar for 5’ and 3’ exon ends, with a marginally smaller contribution of 3’ exon ends (Supplementary Figure 1). This might be linked to a greater fraction of nucleotides at 3’ exon termini being under selection to maintain splice-relevant base-pairing interactions with the neighbouring intron (see below). Inevitably given the area of SNP enrichment, a substantial proportion of boundary-proximal polymorphisms are located in known endonuclease cleavage motifs (17/24=71% of mutations within 20nt of the intron-exon boundary overlap homing endonuclease recognition sites). This might be considered surprising. However, systematic mutagenesis experiments previously demonstrated that many single-nucleotide changes do not perturb target recognition and cleavage (Sargueil *et al*. 1990; Wernette *et al*. 1992). Indeed, the three SNPs recovered here that overlap a previously mutagenized endonuclease cleavage site (split across exon 4 and 5 of *cox1*) had all been tested individually and found to have wild-type cutting efficacies (Sargueil *et al*. 1990).

### SNP density gradients are specifically associated with mobile introns

Previous studies comparing pairs of strains (one with and one without a focal intron) had postulated the presence of mutational hotspots near intron-exon boundaries but lacked quantitative support (Hensgens *et al*. 1983; Zimmer *et al*. 1987; Foury *et al*. 1998). These studies had also speculated that mobility might be a causal factor in elevated nucleotide diversity, or more specifically, that variation was being introduced upon intron gain (Zimmer *et al*. 1987) or loss (Hensgens *et al*. 1983). The more extensive sampling of population genetic variation carried out here reveals that there is no perfect correspondence between polymorphisms and intron presence or absence (an illustrative example is shown Fig 1C), precluding straightforward attribution of novel exonic variation to intron gain or loss. However, we find strong support that mobility in general is key. Although an elevated SNP density is detectable when considering polymorphisms across all *cob/cox1* exon termini, this effect is specifically driven by exon ends that adjoin mobile introns (τ=−0.63, P=2.3E-14, Fig 2A; mobility as defined by previous experimental research, see Methods). Exon ends bordering immobile mitochondrial introns do not show a similar enrichment for SNPs (τ=0.03, P=0.71). Similarly, we find no SNP enrichment in the exonic borders of spliceosomal nuclear introns, which lack the capacity to excise themselves from their host mRNA and do not encode endonucleases or other mobility factors. Rather, exon boundaries are somewhat depleted of SNPs (τ=0.40, P=3.4E-6, Fig 2B). In short, elevated SNP densities at intron-exon boundaries are confined to introns that are both self-splicing and mobile. Further support for a critical role of mobility comes from population genomic analysis of 1135 *Arabidopsis thaliana* accessions (see Methods), whose mitochondrial and chloroplast genomes also harbour self-splicing introns embedded in protein-coding genes.

**Figure 2.**
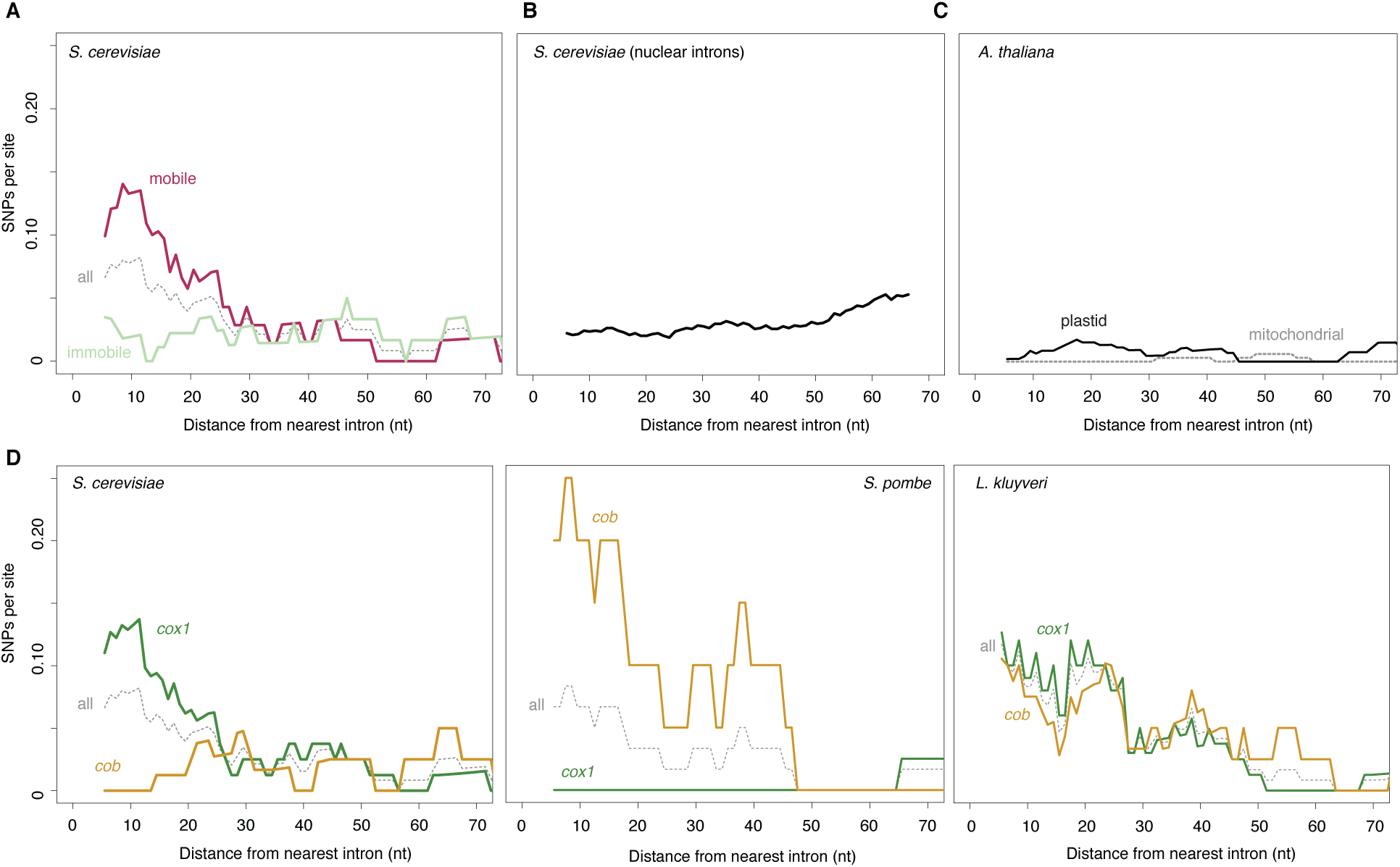
Elevated SNP density at intron-exon boundaries in different genes and species. A SNP density as a function of distance from the nearest intron-exon boundary across *S. cerevisiae cox1/cob* exons experimentally determined to be mobile (see Methods) and the immobile remainder. The grey dotted line (all) indicates the combined mobile/immobile data and corresponds to the data shown inFig 1B. B SNP density as a function of distance from the nearest intron-exon boundary across *S. cerevisiae* exons bordering spliceosomal nuclear introns. C SNP density as a function of distance from the nearest intron-exon boundary for introns in intron-containing mitochondrial (grey) or chloroplast (black) protein-coding genes of *Arabidopsis thaliana*. D SNP density as a function of distance from the nearest intron-exon boundary for introns in the *cob*(yellow) and *cox1* (green) genes of *S. cerevisiae, S.pombe*, and *L. kluyveri*. The grey dotted lines (all) indicate combined *cox1/cob* data.

However, unlike their fungal counterparts, these introns lack open reading frames that encode functional endonuclease, RT or other domains that might mediate mobility and, like those of other land plants, are not mobile as a consequence (Bonen 2008). As predicted under a model where mobility is critically linked to elevated nucleotide diversity, we find no evidence for higher SNP densities near intron-exon boundaries in *A. thaliana* (combined τ=−0.13, P=0.11, Fig 2C), albeit on a background of globally low mutation rates.

In *S. cerevisiae*, most mobile introns are located in *cox1*, with only a single intron in *cob* reported to be mobile in crossing experiments (Lambowitz and Belfort 1993) (Fig 1A). As a consequence, *cox1* exons exhibit SNP enrichment near the intron-exons boundary (τ=−0.67, P=1.59E-15) whereas *cob* exons do not (τ=0.23, P=0.007). To rule out gene-specific factors rather than mobility in the genesis of nucleotide diversity, we examined mitochondrial protein-coding genes from 161 *S. pombe* and 16 *L. kluyveri* strains (see Methods). The *S. pombe* reference genome encodes two introns in *cox1* and a single intron in *cob*. Importantly, the group II *cob* intron alone is known to be mobile (Zimmer *et al*. 1987). In *L. kluyveri*, a recent study found evidence for mobility of both *cox1* and *cob* introns, noting presence/absence polymorphisms for three out of four *cob* and three out of five *cox1* introns (Jung *et al*. 2012). In line with widespread mobility in this species, all introns with the exception of the first *cob* intron encode endonucleases (Friedrich *et al*. 2012). As predicted under a model where gene identity is secondary but mobility plays a pivotal role in nonrandom nucleotide diversity at intron-exon boundaries, we observe SNP density gradients across both *cob* (τ=−0.64, P=2.5E-14, Fig 2D) and *cox1* exons (τ=−0.67, P=1.7E-15) in *L. kluyveri* whereas in *S. pombe* a negative SNP gradient is evident for *cob* (τ=−0.75, P=6.8E-17) but not *cox1* (τ=0.47, P=1.08E-6).

### No evidence for relaxed purifying selection at intron-exon boundaries

A number of evolutionary scenarios might account for elevated exonic nucleotide diversity at sites of intron gain and loss. Importantly, mobility need not be causal: introns might instead be located in areas that are under reduced selective constraint. To investigate whether homing sites might be biased towards regions under lower functional constraint, we considered the ratio of non-synonymous to synonymous changes as an indicator of protein-level selection. In *L. kluyveri*, within 20nt of the intron-exon boundaries of *cox1* and *cob* only six out of 29 SNPs (21%) are non-synonymous, a significant depletion compared to the mutational expectation of ~2/3 (Fisher test P=0.001). Similarly, only one out of eight SNPs (12.5%) in close vicinity of the *S. pombe cob* intron is non-synonymous (Fisher test P=0.12, but note that power here is severely limited by the small number of mutations). In both cases, lower levels of non-synonymous diversity support the notion of strong ongoing protein-level selection at intron-exon boundaries. Interestingly, we find a relatively large number of non-synonymous mutations in *S. cerevisiae cox1* (15/44=34%). However, the ratio is similarly high (5/19=26%) further away from the boundary (Fisher test P=0.77), arguing for a global rather than local, boundary-anchored relaxation of constraint. This observation is broadly consistent with prior evidence for reduced selection on mitochondria in the wake of the whole genome duplication (WGD) event (Jiang *et al*. 2008). We note that, in this regard, *S. cerevisiae* and other post-WGD species might be uniquely informative for assessing mutational forces at work in mitochondrial protein-coding genes.

Further testimony for ongoing purifying selection at the intron-exon boundary comes from scrutinizing exonic residues involved in base-pairing interactions with the neighbouring intron (see Methods). Disruption of proper exon-intron base-pairing is anticipated to impair splicing, with deleterious consequences for the host since splicing is required to reconstitute functional *cob/cox1* reading frames. Although splice-relevant exonic residues are firmly located in the zone of enriched SNP density, we find only a single SNP at a nucleotide position that is involved in exon-intron base-pairing; and even this SNP, a synonymous change (CAC<->CAT) at the 5’ end of *cox1* exon 5, likely preserves base pairing (G-C<->G-T). This observation supports the notion that we do not see higher SNP densities as the result of relaxed selection but rather despite persistent functional constraint. In fact, if excess variation reflects differential mutational input, we are likely to underestimate the true mutational gradient given the added splicing-related constraints in the immediate vicinity of introns.

Finally, to provide a further, complementary layer of evidence that variation in local conservation is not the cause of excess genetic diversity in the vicinity of mobile introns, we make use of the highly conserved nature of ETC proteins across eukaryotes and consider local topologies of constraint in human *cox1* and *cob* (also known as *cytb*). We reasoned that local constraints on protein function and structure should be very similar between the human and yeast orthologs and that, in considering polymorphisms found in the constitutively intronless human orthologs, we circumvent potential circularity in assessing the relationship between introns and local conservation. We therefore charted SNP density across human *cox1* and *cob* as a function of distance from mock splice junctions, placed at orthologous positions in the respective gene (see Methods). We find no evidence for locally relaxed selective constraint for either gene, regardless of whether we assume *S. cerevisiae* or *L. kluyveri* intron positions (Fig 3). Overall, the evidence presented above is inconsistent with locally reduced selection and instead points to a causal contribution of mobility in generating observed diversity patterns.

**Figure 3.**
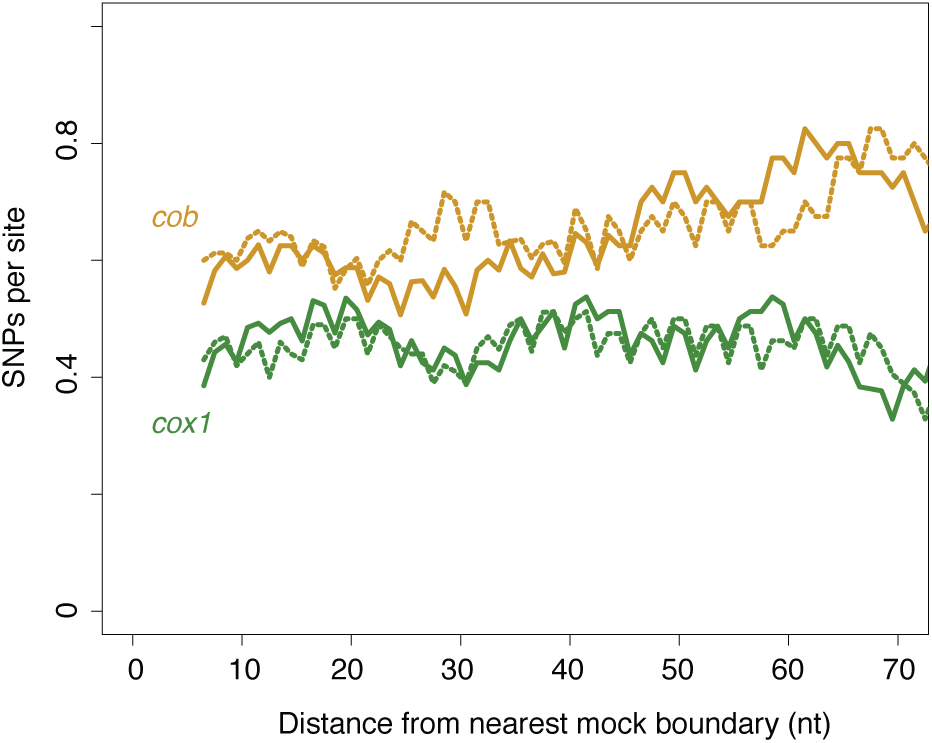
No evidence for locally relaxed purifying selection at intron-exon boundaries. SNP density across human *cob* (yellow) and *cox1* (green) and as a function of mock intron-exon boundaries introduced *in silico* into the human proteins (see main text), based on either intron positions in *S. cerevisiae* (solid lines) or *L. kluyveri* (dotted lines). Note that the 3nt periodicity reflects the fact that the majority of introns occur in the same phase.

### Candidate molecular mechanisms for mobility-associated SNP patterns

Which mobility-associated molecular processes might lead to elevated SNP rates at intron-exon boundaries? One candidate mechanism is gene conversion. Previous studies in yeast, (e.g. Zinn and Butow 1985), and notably also plants (Cho *et al*. 1998; Sanchez-Puerta *et al*. 2011), have provided experimental and comparative genomic support for exon coconversion. After endonuclease activity has introduced a DSB into the intron-free locus, exonucleases resect part of the neighbouring exon and genetic information not previously present is introduced into the exon from the uncleaved repair template. The length of the resected fragment varies, with exonic portions closer to the intron more frequently affected (Mueller *et al*. 1996). However, more frequent conversion of intron-proximal exonic sequence does not by itself explain higher diversity in that region. This is because gene conversion only shuffles pre-existing genetic diversity. If it occurs between genomes in the same recombining population, no additional variation is introduced that would account for a greater incidence of SNPs near exon-intron boundaries. Thus, for exon co-conversion to explain our data, an important additional requirement needs to be met: novel variants must be introduced into the population from the outside. That is, the template for conversion is introduced via horizontal transfer or introgression events. And since we are considering diversity within extant populations, this process has to be ongoing (or at least recent) and pervasive (i.e. affecting several introns). Such a scenario is not necessarily unreasonable given prior findings, notably in plants, of high rates of horizontal transfer of self-splicing introns (Cho *et al*. 1998; Goddard and Burt 1999; Strope *et al*. 2015). However, we were unable to identify likely donors for such putative HGT events, despite a comprehensive survey of NCBI's non-redundant nucleotide database.

An alternative to the HGT plus exonic co-conversion model is that intron insertion sites constitute mutational hotspots, as previously suggested (Hensgens *et al*. 1983; Zimmer *et al*. 1987; Foury *et al*. 1998). There are two major scenarios how mutations might be generated as a side effect of intron mobility. In the first scenario, mutations are introduced when intron-free mRNAs or intron-containing pre-mRNAs are converted into cDNA by a resident error-prone reverse transcriptase, and that cDNA mediates precise intron loss or intron gain, respectively. This process can in principle act in *trans* and impact introns other than those specifically encoding ORFs with RT activity. However, importantly, reverse transcription does not predict a higher mutation load at exon-intron boundaries. In the second scenario, novel variants are produced by mutagenic repair following endonuclease-mediated cleavage. Interestingly, there is some prior evidence – based on studies of the *S. cerevisiae* MAT locus – that DSB repair in the context of endonuclease-mediated cleavage is mutagenic. Yu and Gabriel found deletions in the Z region, which borders the cutting site of the HO endonuclease, in high proportion (2%) of yeast crosses, which were attributed to microhomology-mediated end joining (Yu and Gabriel 2003). Further, studying HR, Hicks and colleagues noted high (1400-fold over spontaneous) rates of predominantly single nucleotide mutations, which they attributed to the action of error-prone polymerases (Hicks *et al*. 2010). Given these prior observations, we naturally examined polymorphisms in the Z1 region of the MAT locus but found it to be perfectly conserved across the 92 *S. cerevisiae* strains analyzed here. The first SNP was found 192nt downstream of the HO cutting site. This high level of conservation is indicative of exceptionally strong nucleotide-level constraint and echoes previous observations of very slow divergence (>96% nucleotide identity) between otherwise well-diverged Saccharomyces *spp*. (Gordon *et al*. 2011). Unfortunately, this precludes the use of natural diversity at the MAT locus as a model to study mutagenic effects.

It is worth considering at this point whether the finding that HO-initiated DSB repair is mutagenic might be specifically reflective of HR following endonuclease-mediated cleavage rather than HR in general. Is it possible that the activity or presence of the endonuclease itself affects the repair process? There have been a number of recent reports that DNA-binding proteins, by associating with a lesion-containing target site, can prevent proper damage surveillance and repair, ultimately leading to a higher incidence of mutations (Reijns *et al*. 2015; Sabarinathan *et al*. 2016; Kaiser *et al*. 2016). Endonucleases, which bind their recognition motifs with high affinity, might elicit similar effects, for example by competing with the repair machinery when DSB repair is being templated by a second intron-free copy of the mitochondrial genome, which is also at risk of being cleaved. Indeed, there is some evidence of repair interference from the self-splicing *td* intron of phage T4, where the intron-encoded I-TevI endonuclease, which cleaves distally to its binding site, remains preferentially associated with one of the free cleavage products and thereby asymmetrically impedes resection (Mueller *et al*. 1996). We posit that endonuclease activity remains an intriguing candidate for the mechanism behind mobility-associated mutagenicity and warrants further investigation. We also suggest that, although HR is generally considered to be relatively error-free, HR in the context of endonuclease-mediated cleavage might follow systematically different repair dynamics – a hypothesis that deserves additional experimental scrutiny given the central role of targetable endonucleases in contemporary biotechnology.

If non-random patterns of genetic diversity are indeed mutational in origin, our findings have important implications for the cost of self-splicing introns, which have generally been considered as relatively cost-free given their ability to efficiently remove themselves from their host genes (Werren 2011). Our results would suggest instead that these introns impose a mutational load on their host genes in addition to potential physiological costs such as the energy and time expended in the splicing process. In as far as this mutational load affects fitness, the mutagenic effect might co-determine what constitutes evolutionarily sustainable insertion sites. That is, long-term safe havens for self-splicing introns may be limited to regions within genes that exhibit sufficient functional constraint so that the host cannot mutate away the cleavage site but are also robust enough to tolerate mutations introduced with some regularity by mutagenic activity. If, on the other hand, exonic co-conversion is responsible for intron-proximal SNP gradients, our results strongly argue for ongoing crossspecies transfer of mobile introns in extant yeast populations on a previously unrecognized scale.

## Methods

### Data acquisition and identification of polymorphic sites

We obtained the sequences of 93 *S. cerevisiae* mitochondrial genomes, originating from a recent high-coverage re-sequencing effort (Strope *et al*. 2015), from John Wolters (Binghamton U.). Baiting BLAST searches (blastn, E value < 1E-9) with the terminal exons of *cob* and *cox1* from the reference S288C genome, we identified unique full-length *cob* and *cox1* genes in all strains, capturing both coding exons and intervening introns. We aligned the 94 sequences (93 plus the S288C reference genome) for each gene together with 50nt of upstream/downstream flanking DNA using MUSCLE v. 3.8.31 (Edgar 2004) and manually surveyed the alignment around intron-exon boundaries for alignment errors or conspicuous outliers. As a consequence of this manual inspection, we conservatively excluded strain YJM1250, which exhibits an unusual multi-nucleotide difference at the 5’ end of exon 6, which would have further exacerbated the SNP density gradient reported here. We also excluded strain YJM1399, whose mitochondrial genome was previously found to be more closely related to *S. paradoxus* (Wolters *et al*. 2015). Polymorphic sites in *cob* and *cox1* were therefore identified from the alignment of 92 sequences. In inferring distances to the nearest intron-exon boundary, we only considered introns present in the reference genome. This is conservative since residues that are inferred to be exon-internal might in fact be close to an intron present in the population but not the reference genome, thus overestimating mutations internal to the exon.

For the analysis of the MAT locus and nuclear spliceosomal introns, chromosomes for the 91 re-sequenced yeast genomes and S228C were downloaded via Batch Entrez based on their GenBank accessions in (Strope *et al*. 2015) and the Saccharomyces Genome Database (http://www.yeastgenome.org), respectively. Identification and alignment of the MATα (which is reported in all genome assemblies in favour of MATa) and Z1 regions based on the S288C annotation was straightforward given the exceptional conservations levels reported above. Genes containing nuclear spliceosomal introns in S228C were identified based on GenBank annotations. The terminal exons of these genes (required minimum length >1nt) were then blasted against the remaining genomes (blastn, E value < 1E-9). Homologous sequences were extracted for cases where both terminal exons were at least 70nt long, identified as the only hits in the BLAST query, located on the same chromosome and strand and less than 3kb apart (covering the empirical intron-containing gene length distribution in *S. cerevisiae)*. Homologous sequences recovered in at least 80 strains were then aligned using MUSCLE with default parameters.

We obtained *cox1* and *cob* coding sequences for 18 *L. kluyveri* strains from Paul Jung (U. of Luxembourg). Intron positions in these strains were taken from Supplementary Figure 2 of (Jung *et al*. 2012). Following alignment of these sequences (using the same parameters employed for *S. cerevisiae)* and subsequent manual inspection, we conservatively excluded strains CBS10367 and CBS10368 because of conspicuous divergence at the 5’ end of exon 4. In inferring distances to the nearest intron-exon boundary, we considered all intron insertion sites observed across the 18 *L. kluyveri* strains by (Jung *et al*. 2012).

We obtained variant calls across 161 *S. pombe* strains (Jeffares *et al*. 2015) from Daniel Jeffares (available at https://danielieffares.com/data/). In the absence of high-quality *de novo* mitochondrial assemblies for these strains and unknown intron presence/absence variability, we only considered introns present in the *S. pombe* reference genome when inferring distances to the nearest intron-exon boundary.

Variant calls across 1135 *A. thaliana* chloroplast and mitochondrial genomes were obtained from the 1001 Genomes Project (Cao *et al*. 2011) (via Ashley Farlow). In the absence of *de novo* mitochondrial/chloroplast assemblies for these strains, we only considered introns present in the *A. thaliana* TAIR10 reference genome when inferring distances to the nearest intron-exon boundary.

Human *cob* and *cox1* coding sequences and associated polymorphism data were obtained from the Mitomap database (Lott *et al*. 2013). Mock intron-exon boundaries in the human sequences were placed at orthologous positions as identified from human-*S. cerevisiae*-*L. kluyveri* alignments for *cox1* and *cob*.

### Calculation of SNP densities and assessment of overlap with functional motifs

For all species, SNP density at a given distance from the boundary is calculated across all pertinent exons as the number of polymorphic sites per total sites at that distance. Correlations were calculated across a 70nt window from the boundary, which captures the regions of elevated and plateauing SNP density in all species. Further extending this region is not beneficial since increasingly fewer exons contribute to specific positions.

We use structural information provided by the Group I Intron Sequence and Structure Database [GISSD, (Zhou *et al*. 2007)] and the Zimmerly lab (http://www.fp.ucalgary.ca/group2introns/) to assess the degree to which polymorphic exonic sites were involved in pairing to intronic residues as part of the self-splicing process. Overlap with endonuclease cleavage sites was assessed based on cleavage motifs defined in REBASE v608 (Roberts *et al*. 2010). Experimentally mobile *S. cerevisiae* introns were defined as in (Lambowitz and Belfort 1993). Polymorphisms were classified into synonymous and non-synonymous according to the yeast mitochondrial code (NCBI transl_table=3) for *S. cerevisiae* and *L. kluyveri* and the standard code (NCBI transl_table=1) for *S. pombe*.

## Acknowledgments

We thank John Wolters, Paul Jung, Daniel Jeffares, Jonathan Flowers, Ashley Farlow and Detlef Weigel for sharing sequence data and alignments, and Peter Sarkies and Anita Krisko for comments on the manuscript. TW is the recipient of intramural funding from the UK Medical Research Council (MRC) and an Imperial College Junior Research Fellowship.

## Author contributions

JR and TW performed data analysis. TW conceived the study, designed analyses, and wrote the manuscript.

